# Independent mechanisms underlie the protective effect of dietary polyunsaturated fatty acid supplementation and Gα_z_ deficiency on the early type 1 diabetes phenotype of Non-obese diabetic (NOD) mice

**DOI:** 10.1101/2021.03.13.435254

**Authors:** Rachel J. Fenske, Haley N. Wienkes, Darby C. Peter, Michael D. Schaid, Andrea Pennati, Jacques Galipeau, Michelle E. Kimple

## Abstract

Non-obese diabetic (NOD) mice deficient in G_z_ alpha subunit (Gα_z_) are protected from developing hyperglycemia, even with early islet insulitis similar to wild-type mice. Similarly, wild-type (WT) NOD mice are protected from glucose intolerance when fed a diet enriched in eicosapentaneoic acid (EPA). In the beta-cell, Prostaglandin EP3 receptor (EP3), whose primary endogenous ligand is the arachidonic acid (AA) metabolite, prostaglandin E_2_, is specifically coupled to Gα_z_. In this work, we tested whether dietary EPA supplementation, thereby reducing systemic PGE_2_ levels, would complement Gα_z_ loss in the NOD mouse model. WT and Gα_z_-null NOD mice were fed an AA-enriched diet, EPA-enriched diet, or control diet upon weaning. After 12 weeks of diet feeding, glucose tolerance tests were performed and pancreatic islets and whole pancreas collected for *ex vivo* analyses, with the longer-term effect of an EPA-enriched diet on splenic T-cell populations quantified via flow cytometry. Our results reveal a polyunsaturated fatty acid-enriched diet, whether AA or EPA, improves wild-type NOD glucose tolerance by the same magnitude as Gα_z_ loss, but through almost completely different physiological and cellular mechanisms. Our results shed critical light on future research into novel pharmacological and dietary adjuvant therapies for T1D.

## Introduction

Type 1 diabetes (T1D) is a chronic autoimmune disease that affects individuals of all ages, but has historically been diagnosed in childhood. Over the past decades, the prevalence of T1D and other autoimmune diseases have been increasing. In the United States, the incidence of new onset T1D is 10-25 in every 100,000 births and globally, the prevalence of T1D in individuals under the age of 18 is over 500,000 (1). While exogenous insulin therapy allows individuals with T1D to lead relatively normal lives, the management of T1D can still be quite difficult. Insulin dosing, exercise, stress, illness and a host of other factors can complicate blood glucose management. New treatments are desperately needed.

It is now well-accepted T1D onset and progression involve both immune system dysregulation and underlying β-cell dysfunction. Although an ever-increasing number of immune-modulatory therapies have entered phase 3 clinical trials, none have been fully successful in achieving goal outcomes. Research into β-cell-centric strategies to cure or treat T1D have focused on three general areas: fine-tuning the pharmacological properties of exogenous insulin and its dosing mechanisms to better replicate the endogenous state; improving islet transplantation outcomes and developing and validating β-cell replacement strategies; and identifying therapeutic targets to stimulate the regeneration of existing β-cells and improve their function *in vivo*. A dietary intervention could be an effective complement to one or more classes of interventions. Unfortunately, there is a dearth of conclusive evidence-based dietary recommendations for individuals with T1D (2), and dietary interventions for T1D management have come under recent and intense scrutiny due to the popularity of fad diets, particularly in this high-risk population (3).

Linoleic acid (an omega-6 PUFA) and alpha-linolenic acid (an omega-3 PUFA) are converted *in vivo* to arachidonic acid (AA) and eicosapentaneoic acid (EPA), respectively, through desaturation and elongation. In addition to being biologically-active compounds in their own right, AA-derived PGE_2_ and EPA-derived PGE_3_ stimulate a family of prostanoid receptors that have been well-characterized to have important roles in metabolically active tissues, including islets, adipose tissue, and immune cells. Prostaglandin EP3 receptor (EP3) has long been a focus of research in both the T1D and T2D fields. Our lab has shown β-cell EP3 can couple to the unique inhibitory G protein alpha subunit, Gα_z_, to block β-cell function, replication, and survival (4–6). Mice deficient in Gα_z_ are protected from developing T1D-like hyperglycemia, whether instigated by the β-cell toxin, streptozotocin, or in the non-obese diabetic (NOD) background (7–9). Based on average human and murine diets, AA is much more abundant in cellular membranes than EPA, including the pancreatic islets (9). While the current body of literature clearly supports an essential role of PGE_2_-mediated signaling through EP3 in the β-cell dysfunction of T2D, there is evidence this mechanism is conserved in T1D.

A preliminary study from our laboratory confirmed feeding NOD mice a semi-purified diet enriched with a physiologically-relevant concentration of EPA improved their glucose tolerance: an early step in the progression to T1D. In this work, we aimed to determine the specific mechanisms behind this protection and whether they were dependent on Gα_z_. Wild-type (WT) and Gα_z_-null (Gα_z_KO) NOD mice were fed an EPA- or AA-enriched diet, or matched control diet, upon weaning. Oral glucose tolerance tests were used as a measure of progression to T1D, and immune phenotype and β-cell function were quantified at the systemic and islet levels. Our results shed light on the specific effects of EPA, AA, and their metabolites, on the organismal and cellular mechanisms underlying T1D pathophysiology and the requirement of β-cell Gα_z_ in mediating these effects.

## Materials and Methods

### Animals

Congenic NOD/ShiLtJ (NOD) containing a genomic insertion of a pGKneor cassette 160 base-pairs downstream of the translation start site of the Gα_z_ gene (gene symbol: *Gnaz*) have been described, validated, and characterized previously (8). Breeding colonies were housed in a limited access, pathogen-free facility (UW Madison Biomedical Research Model Services Mouse Breeding Core) where all cages, enrichment, and water were sterilized prior to use. Mice were maintained on a 12 h light/dark cycle with ad libitum access to water and irradiated breeder chow (Teklad 2919). Gα_z_-null (Gα_z_KO) and wild-type (WT) NOD mice were generated by heterozygous matings to produce littermate pairs as previously described (8). Upon weaning, mice were transferred to an investigator-accessible facility and housed two per cage of mixed genotypes with ad libitum access to a modified AIN-93G base diet, with or without the addition of 2 g/kg EPA or AA, as described previously (9). Briefly, to limit endogenous AA and EPA production from shorter chain omega-6 and omega-3 PUFAs, coconut oil containing primarily short chain saturated fatty acids replaced soybean oil as the predominant fat source in the modified AIN-93G base diet.

Female NOD body weight, food consumption (calculated by subtracting pellet mass remaining from pellet mass initially provided), and random-fed blood glucose were measured weekly from weaning, for 12 weeks, until final phenotyping and terminal tissue collection. Blood glucose measurements were recorded using an AlphaTRAK blood glucose meter and rat/mouse-specific test strips. A contemporaneous cohort of NOD male mice was used to collect pancreatic islets after 12 weeks of diet feeding, and a separate cohort of female WT and Gα_z_KO NOD were housed until 28-29 weeks of age before euthanasia and splenic T-cell isolation.

All protocols were approved by the Institutional Animal Care and Use Committees (IACUCs) of the University of Wisconsin-Madison and the William S. Middleton Memorial Veterans Hospital and all facilities were accredited by the Association for Assessment and Accreditation of Laboratory Animal Care (AAALAC). All animals were treated in accordance with the standards set forth by the National Institutes of Health Office of Animal Care and Use.

### Oral glucose tolerance tests and quantification of plasma insulin and PGE2 metabolite (PGEM) levels

16-17-week old NOD females were fasted for 4 to 6 hours before glucose tolerance testing as previously described (6,9). Briefly, glucose was bolused by oral gavage at a dose of 2 g/kg body weight. Blood glucose measurements were taken immediately before gavage and at 5, 15, 30, 60, and 120 minutes following gavage. EDTA-anticoagulated blood was collected using a lateral tail nick at baseline and 5 minutes after gavage for plasma insulin measurements (Ultra Sensitive Mouse Insulin ELISA; Crystal Chem; catalog #90080). Prior to euthanasia for terminal tissue collection, blood was collected by retroorbital puncture into heparin-coated microcapillary tubes for downstream analysis of Prostaglandin E_2_ metabolite (PGEM) (Prostaglandin E Metabolite ELISA kit; Cayman Chemical; catalog #514531). ELISA assays were performed according to the manufacturers’ protocols, as previously described (4,10).

### Quantification of insulitis severity and β-cell fractional pancreas using formalin-fixed, paraffin-embedded slide sections

Whole pancreas was dissected and fixed in 10% formalin for 48 hours followed by storage in 70% ethanol (4°C) until paraffin embedding. 10-micron serial sections were cut on positively charged slides. Immune infiltration of the β-cell was determined using standard eosin (Polysciences Cat #17269) and hematoxylin (Sigma, GHS280) staining. Following staining, quantification of immune infiltration was accomplished using a numerical scoring system, as previously described (8,9). Every islet in a single section was counted and two 10-micron sections, separated by at least 200 microns, were scored and averaged per mouse.

Beta-cell fractional area was quantified on serial slide sections as previously described (4,6,7,9,11,12). Briefly, pancreas sections were post-fixed, quenched of peroxidase activity with 3% H_2_O_2_, and immunohistochemically labeled using guinea pig anti-insulin primary antibody (Dako A056401-2), diluted 1:500 in antibody diluent, and co-stained with hematoxylin (Sigma, GHS280). Slides were imaged using an Evos^®^ automated pan-and-stich microscope at 10x magnification. β-cell fractional area was determined by quantifying the percent of insulin-positive pancreas area as a total of the full pancreas area for each section. Two sections were quantified per mouse and averaged for each biological replicate. Images were analyzed using ImageJ (64-bit) software (National Institutes of Health, Bethesda, MD) with shading correction.

### Islet isolation and glucose-stimulated insulin secretion (GSIS) assays

Mice were euthanized using 2,2,2-tribromoethanol (Sigma; #T48402) anesthesia followed by cervical dislocation. For *ex vivo* islet isolation, collagenase perfusion of the pancreas to isolate pancreatic islets for analysis was performed as previously described (13). On the day of isolation, islets were picked into islet medium [RPMI 1640 (Gibco #11879020) containing 11.1 mM glucose (Fisher Scientific;#D16), 10% heat-inactivated fetal bovine serum (Sigma-Aldrich; #12306C), and 1% HEPES(Sigma; #H4034) and penicillin/streptomycin (Gibco; #15070-063)] and recovered overnight prior to GSIS assays. GSIS treatments included 1.7 mM glucose, 16.7 mM glucose, or 16.7 mM glucose plus 10 nM sulprostone (Sigma-Aldrich; catalog #S8692), and assays were performed and analyzed essentially as previously described (4,6–10,12,14,15).

### Relative gene expression analyses using quantitative real-time polymerase chain reaction (qPCR) of islet cDNA samples

Relative gene expression was quantified by qPCR using cDNA samples from isolated pancreatic islets essentially as previously described (6–10,14,16). Briefly, isolated islets were hand-picked into islet media and allowed to recover 24 hours as described above. Islets were then picked and washed in PBS before lysis and isolation of whole RNA using the Qiagen RNeasy^®^ Kit (Qiagen; #74106) according to the manufacturer’s instructions. Complementary DNA (cDNA) was then generated with random hexamers (High-Capacity cDNA Reverse Transcription Kit, Applied Biosystems^®^; #4368813), and quantitative reverse transcription polymerase chain reaction (qRT-PCR) was performed using SYBR green (Roche; #04913914001). Quantification of messenger RNA (mRNA) expression was normalized to β-actin cycle times within each sample. Primer sequences are available upon request.

### Spleen dissection, cell Isolation, and flow cytometry procedures

Spleens were harvested postmortem from 28-29-week-old female WT or Gα_z_KO mice and collected in RPMI medium (Lonza). Cell isolation and flow cytometry procedures were conducted as previously described (17). Briefly, polarized cells were washed and stained with Ghostred for 30 min at 4 °C. Surface staining with anti-CD3, anti-CD4 and anti-CD25 was carried out at 4 °C for 30 min. For intracellular staining, polarized cells were washed, placed in fresh medium, and cultured with 50 ng/ml PMA and 500 ng/ml ionomycin at 37 °C for 5 h. Brefeldin A (10 μg/mL) was added for the last 3 h of incubation. After staining with surface markers, cells were treated with freshly prepared fixation/permeabilization buffer (Invitrogen) and incubated at 4 °C for 30 min. After washing with 1× permeabilization buffer (Invitrogen), intracellular staining was performed at 4 °C for 30 min using anti-IFN-*γ*, anti-IL-17 and anti-FoxP3. All antibodies were from Invitrogen. AttuneNxT was used for data collection, and FCS Express software was used for data analysis.

### Statistical analysis

Data are expressed as mean ± standard error of the mean (SEM) unless otherwise noted. Statistical analyses were performed as described in each of the figure legends using GraphPad Prism version 9.0 (GraphPad Software, San Diego, CA).

### Data and Resource Availability

The datasets generated during and/or analyzed during the current study are available from the corresponding author upon reasonable request.

## Results

### Euglycemic Gα_z_-null female NOD mice are more glucose tolerant than WT females, and a PUFA-enriched diet improves glucose tolerance in both groups

In our NOD colony, female mice do not start becoming hyperglycemic until approximately 20 weeks of age, with full phenotype penetrance (~75%) between 30-35 weeks of age (8). In order to identify mechanisms associated with the T1D pathogenesis and compare among groups without the confounding effects of uncontrolled hyperglycemia, cohorts of WT and Gα_z_-null (Gα_z_KO) females were fed an AA-enriched, EPA-enriched or semi-purified control diet from weaning (between 3-4 weeks of age) for 12 weeks, until 16-17 weeks of age. Body weights, random-fed blood glucose levels, and food intake were recorded weekly, with no significant differences observed by genotype or diet (**Figure 1A**). Circulating PGE_2_ metabolite (PGEM) levels were approximately 30% increased after 12 weeks of feeding an AA-enriched diet and approximately 80% decreased after 12 weeks of feeding an EPA-enriched diet, both as compared to control, with no differences by genotype (**Figure 1B**). Having validated our model, end-of-study phenotyping and terminal tissue collection was performed.

**Figure 1:**
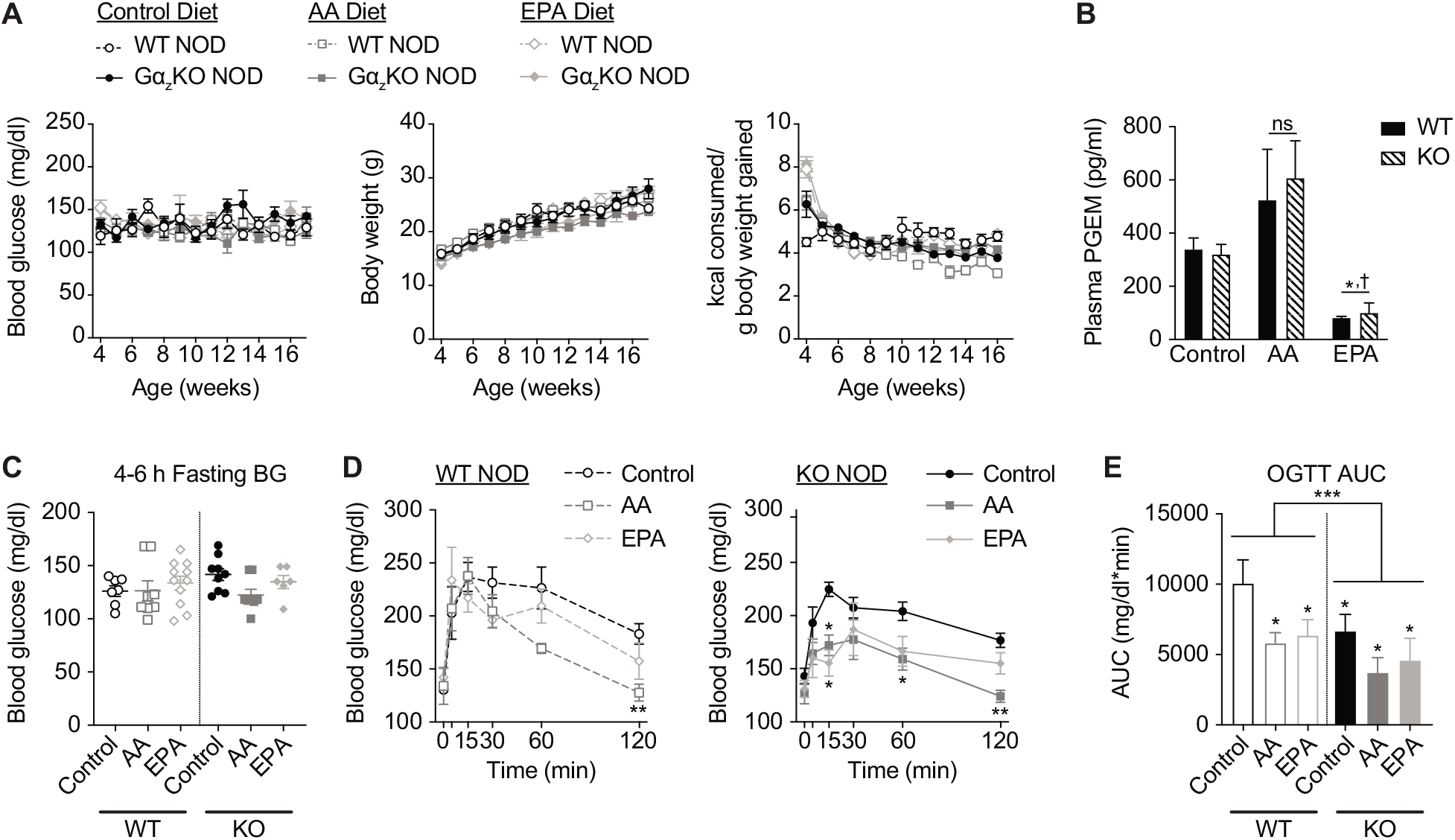
PUFA diet feeding in WT mice and Gα_z_ loss both improve glucose tolerance of female NOD mice. A: Random-fed blood glucose levels (left) body weight (middle) and food consumption (right) of female NOD mice fed a PUFA-enriched diet or control diet for 12 weeks, from weaning (3-4 weeks of age) until 16-17 weeks of age. N=8-11 per group. B: Plasma PGE_2_ metabolite (PGEM) levels after 12 weeks of dietary intervention. N=3-6 per group. *, p < 0.05 vs. Control and †, p < 0.05 vs. AA. C: 4-6 h fasting blood glucose levels after 12 weeks of dietary intervention. N=6-11 per group. D: Blood glucose excursions during oral glucose tolerance tests (OGTTs) after gavage with 2 mg/kg glucose in WT NOD females (left) and Gα_z_KO NOD females (right). *, p < 0.05 and **, p < 0.01 vs. Control. E: Area-under-the-curve (AUC) analyses of the data shown in (D). *, p < 0.05 vs. WT Control. In all panels, data are represented as mean ± SEM, and data were compared by two-way ANOVA with Holm-Sidak test post-hoc to correct for multiple comparisons. In (A) and (D), a repeated measures analysis was used. If a comparison is not shown, it was not statistically significant.

Glucose intolerance precedes overt hyperglycemia in both T1D mice and humans (18). No significant differences were observed in 4-6 h fasting plasma glucose levels among any of the groups, by diet or genotype (**Figure 1C**). In WT NOD females, mice fed either an EPA- or AA-enriched diet had lower mean blood glucose levels at the 30, 60, and 120 min time point (with the AA group being statistically-significant at 120 min) (**Figure 1D, left**). Female Gα_z_-null NOD mice had glucose tolerance excursions similar to those of WT PUFA diet-fed mice, and both EPA diet-fed and AA diet-fed Gα_z_KO females had further reduced blood glucose levels at 15, 30, 60, and 120 min (**Figure 1D, right**). Calculating the mean OGTT AUCs reveals both PUFA diet feeding and Gα_z_ deficiency reduced body burden of glucose nearly 2-fold during the 120 min OGTTs, with the lowest OGTT AUCs being in PUFA-diet fed Gα_z_KO mice (although the latter results were not statistically different than Gα_z_KO alone) (**Figure 1E)**.

### Gα_z_ loss, but not a PUFA-enriched diet, significantly reduces islet insulitis and increases β-cell fractional area

Islet insulitis was quantified from hematoxylin and eosin-stained serial pancreas sections using a validated scoring system, as previously described (8). Control diet-fed Gα_z_KO NOD females had significantly reduced insulitis as compared to WT controls, and this reduction was not impacted by PUFA diet feeding, whether represented as the percentage of total islets with each insulitis severity score (**Figure 2A**) or as the mean islet infiltration score (**Figure 2A**). Notably, about 50% of Gα_z_KO islets showed no evidence of immune cell infiltration (Score 0), with another 25% of islets showing peri-insulitis only (Score 1): a dramatic improvement as compared to WT NOD islets, where 50% were partially (scores 2 and 3) or completely (score 5) infiltrated by immune cells.

**Figure 2.**
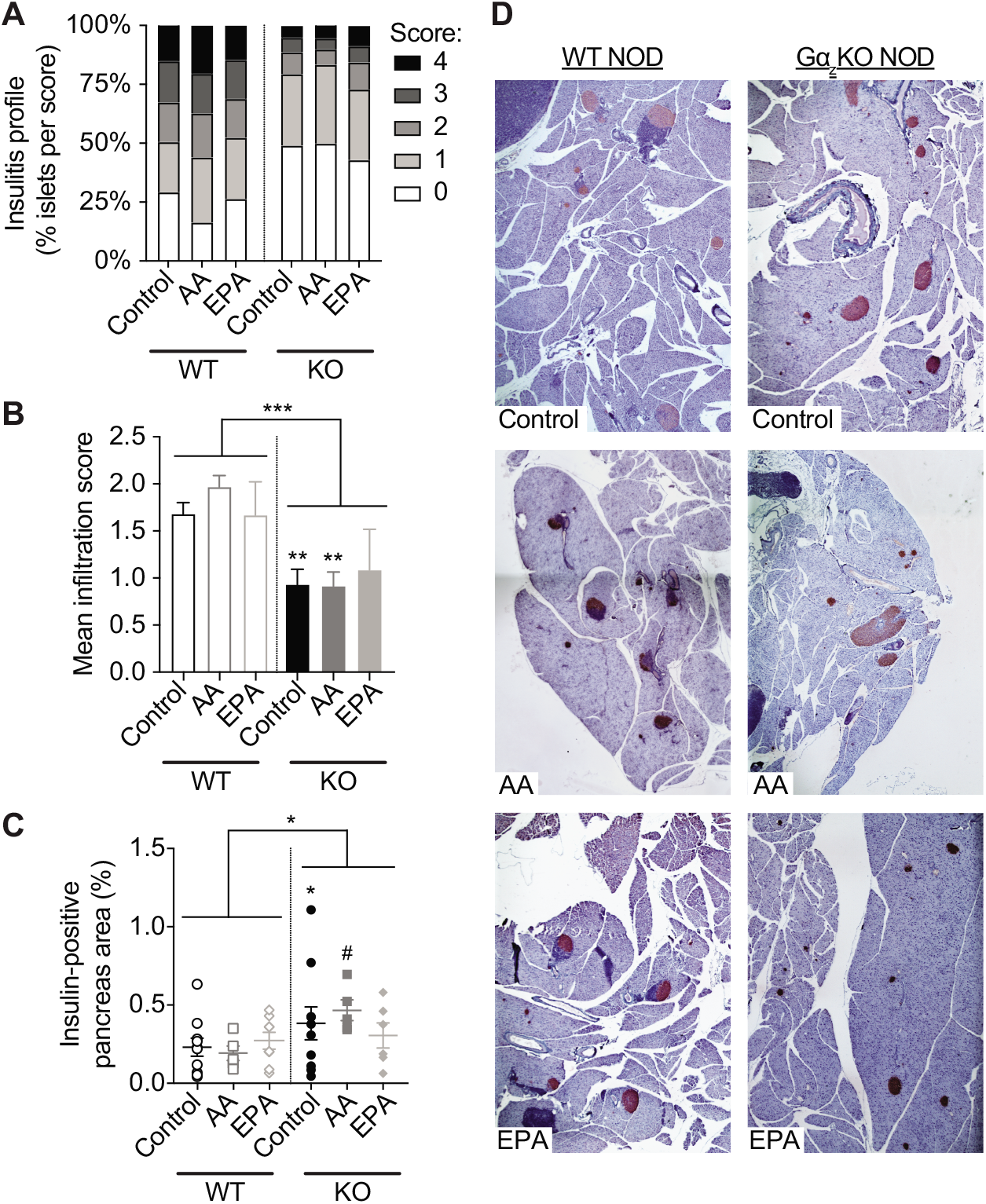
Gα_z_ loss alone significantly reduces islet insulitis and augments β-cell fractional area in 16-17-week old female NOD mice independent of a PUFA dietary intervention. A,B: Degree of islet insulitis as quantified by (A) the percent of islets with each insulitis severity score (B) or the mean islet infiltration score. In (B), data represent mean ± SEM and were compared by two-way ANOVA with Holm-Sidak test post-hoc to correct for multiple comparisons. ***, p < 0.001 for effect of genotype. **, p < 0.05 vs. WT Control. N=6-10 per group. C: Insulin-positive pancreas area as quantified by insulin immunohistochemistry of hematoxylin-counterstained pancreas slide sections. Data represent mean ± SEM were compared by two-way ANOVA with Holm-Sidak test post-hoc to correct for multiple comparisons. *, p < 0.05 for effect of genotype. #, p = 0.10 vs. WT AA diet-fed mice. N=6-10 per group. D: Representative images for the analysis shown in (C).

In order to determine whether differences in islet insulitis correlated with preserved functional β-cell mass, formalin-fixed, paraffin-embedded pancreas sections were subjected to insulin immunohistochemistry with hematoxylin counter-stain and images analyzed in ImageJ. Two-way ANOVA analysis indicated a significant effect of genotype on β-cell fractional area (**Figure 2C**), consistent with previous findings in chow-fed WT or Gα_z_KO NOD mice (8). PUFA diet feeding had no effect on β-cell fractional area in either cohort: although the mean insulin-positive pancreas area of AA-fed Gα_z_ KO mice was higher than control-diet fed, this was not statistically-significant (#, p = 0.10) (**Figure 2C, right**). (Representative images for insulin-positive pancreas area scoring, which also show examples of islet immune infiltration, are found in **Figure 2D**).

### Gα_z_ loss and a PUFA-enriched diet have divergent effects on islet and systemic immune phenotypes

Although the penetrance of hyperglycemia development in male mice is much lower than females (~15% in our colony), when male NOD mice are fed standard chow, the insulitis severity and phenotype with Gα_z_ loss are nearly identical to those of female mice (8). Therefore, in this and our previous work, islets from male NOD mice were also used in gene expression and insulin secretion assays. Interleukin 1 beta (IL-1β) and interferon gamma (IFN*γ*) were both lower in islets from Gα_z_KO mice, with little effect of a PUFA-enriched diet (**Figure 3A,B**). CXCL10 expression was lower in islets from Gα_z_KO mice, but was further reduced by a PUFA-enriched diet (**Figure 3C**). IL-11 expression, on the other hand, was significantly elevated by both a PUFA-enriched diet and loss of Gα_z_, with no additive effect of the two (**Figure 3D**). Finally, NFkB expression was relatively unaffected by either genotype or diet, save for a small but statistically-significant decrease in the EPA diet-fed Gα_z_KO mice as compared to control (**Figure 3E**).

**Figure 3:**
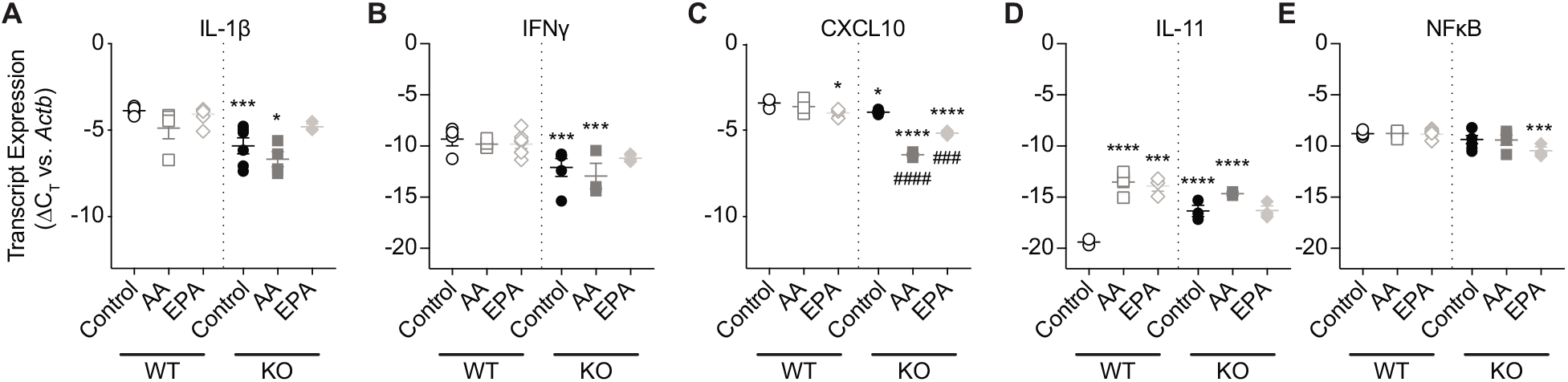
Islet expression of inflammatory cytokines is both independently and coordinately altered by PUFA-enriched diet feeding and Gα_z_ loss. A-E: Results of quantitative PCR (qPCR) analysis of male NOD islet cDNA samples using primers specific for (A) interleukin 1-beta (IL-1β), (B) interferon gamma (IFNɣ), (C) CXCL10, (D) interleukin 11 (IL-11) and (E) NFκB. In all panels, data represent mean ± SEM. Data for each transcript were compared between genotypes by 2-way ANOVA and within genotypes by multiple T-tests, both with Holm-Sidak test post-hoc to correct for multiple comparisons. *p<0.05 vs WT Control. ***p<0.001 vs WT Control. ****p<0.0001 vs WT Control. ^###^p<0.001 vs Gα_z_KO Control. ^####^p<0.0001 vs Gα_z_KO Control. N=3-6 per group.

Because of the known role of prostaglandin signaling in immune cell function, the lack of effect of a PUFA-enriched diet on islet insulitis and, in most cases, islet cytokine expression was unexpected. To confirm whether either diet or genotype affected the systemic immune phenotype of NOD mice long-term, splenic cells were isolated from WT and Gα_z_KO NOD females fed the control diet or EPA-enriched diet from weaning until 28-29 weeks of age. Phenotypic analysis of T cells was carried out based on the expression of IFNɣ (Th1 cells), IL-17 (Th17 cells), and CD25/FoxP3 (classical regulatory T cells, Tregs) **Figure 4A**). Regardless of genotype, splenic T-cells populations shifted as a result of EPA-enriched feeding, with a robust increase in Th1 cells and Tregs, and no change in Th17 population (**Figure 4B**).

**Figure 4.**
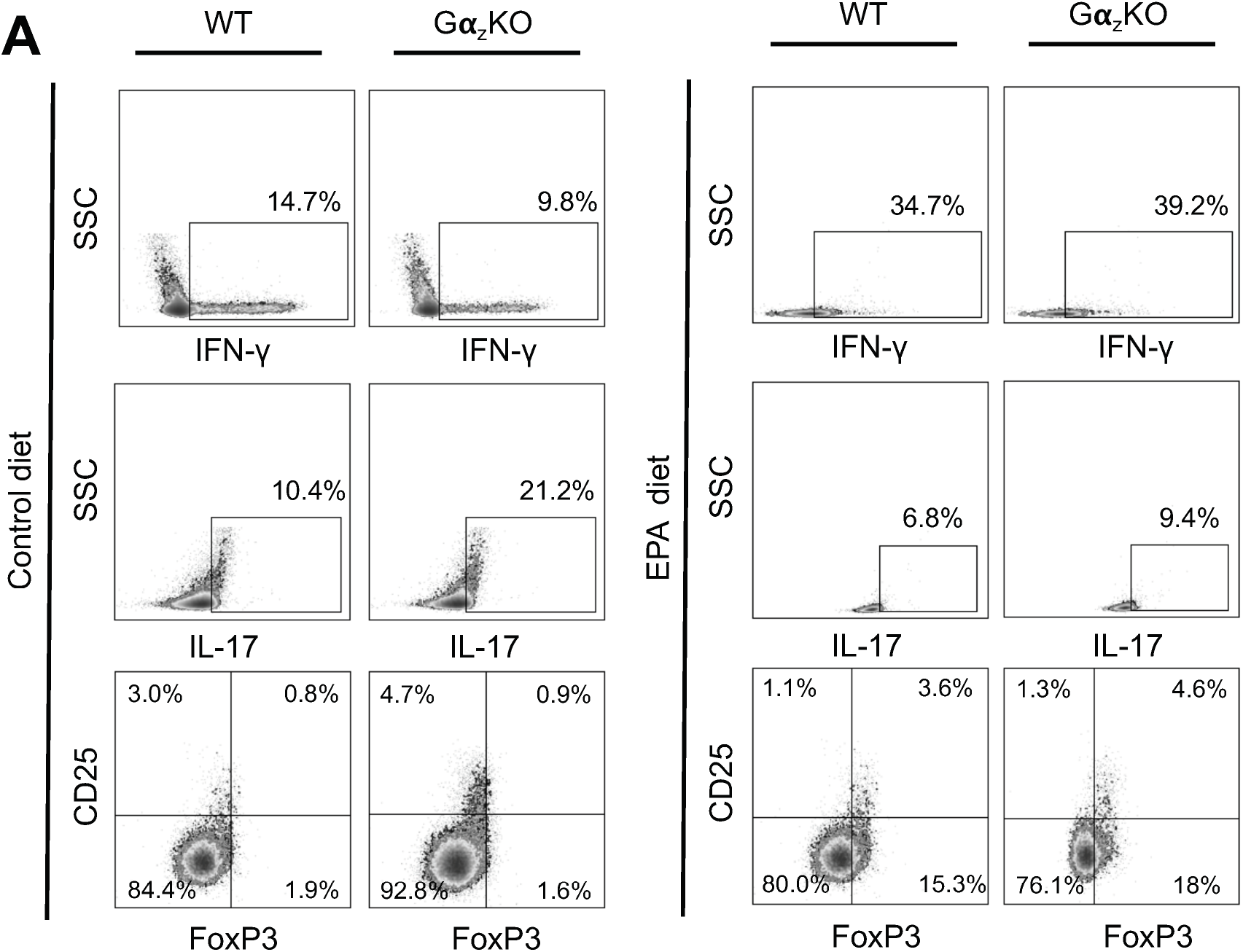
An EPA-enriched diet from weaning reduces alters splenic T-cell populations in 28-29-week-old female NOD mice. Splenic T-cells were extracted from WT and Gα_z_KO NOD females and analyzed by flow cytometry. A: Flow cytometry profiles of control diet-fed mice (left) and EPA diet-fed mice (right). B: Th1, Th17, and Treg populations were compared between group. Data represent mean ± SEM and were compared by 2-way ANOVA with Tukey test post-hoc to correct for multiple comparisons. **p<0.01 vs Control, by genotype. ****p<0.0001 vs Control, by genotype. N=3-4 per group. If a comparison is not shown, it was not statistically-significant.

### *Islets from Gα_z_-null NOD mice secrete more insulin in response to glucose* in vivo *and* ex vivo, *consistent with elevated expression of EP3 splice variants*

Improved glucose tolerance of PUFA diet-fed and Gα_z_KO mice suggested the possibility of an effect on β-cell function. Blood was collected during the 0 min and 5 min OGTT time points for plasma insulin ELISA, but not all blood samples generated enough plasma for the assay. Ultimately, matching 0 and 5 min timepoint plasma insulin measurements were obtained from N=4 control diet-fed mice, N=1 mice AA diet-fed mice, and N=2 EPA diet-fed mice per genotype. Plotting data by diet group revealed Gα_z_KO NOD females secreted, on average, more insulin at baseline and 5 min after oral glucose challenge, with no apparent differences by diet (**Figure 5A**). Therefore, data within genotypes were combined by diet to increase the statistical power of our analysis. 4-6 h fasting plasma insulin levels were significantly elevated in Gα_z_KO vs. WT mice, a finding that held true whether data from AA diet-fed mice were included (*) or excluded (†) in the analysis (**Figure 5B, left**). The same trend existed for 5 min blood glucose values, although the comparison was not statistically significant (**Figure 5B, right**). The fold-stimulation of insulin 5 min after glucose challenge vs. baseline was nearly identical between the genotypes, indicating differences in baseline plasma insulin values as the primary drivers of the phenotype (**Figure 5C**). Plotting the 4-6 h fasting plasma insulin values vs. the corresponding 4-6 h blood glucose values shown in Figure 1C revealed a strong, linear correlation in Gα_z_KO NOD mice, but not WT (**Figure 5D**). Grouping data by genotype revealed Gα_z_KO NOD females had significantly higher 4-6 h fasting insulin:glucose ratios than WT females, regardless of whether data from AA diet-fed mice were included (*) or excluded (†) in the analysis (**Figure 5E**).

**Figure 5.**
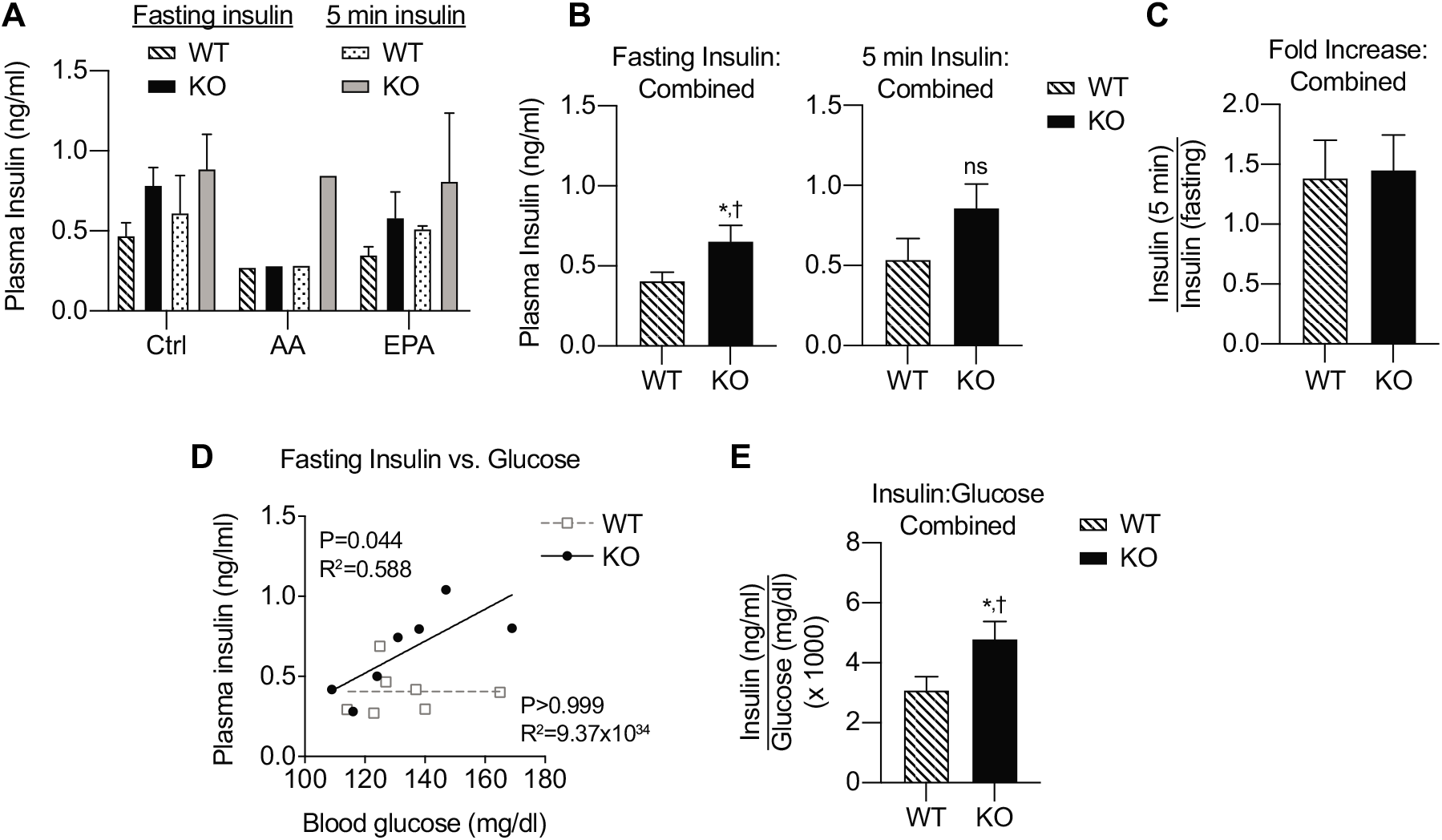
Female Gα_z_KO NOD mice have higher baseline plasma insulin levels for corresponding blood glucose levels than WT controls during the OGTTs shown in Figure 1. A-C: 4-6 h fasting plasma insulin levels or plasma insulin levels 5 min after oral glucose challenge with (A) each group represented separately, (B) results combined by genotype, or (C) 5 min plasma insulin values normalized to their corresponding fasting level. In all panels, data represent mean ± SEM. No statistical analysis was performed in (A). In (B) and (C), data were compared by twoway T test. * or †, p < 0.05 vs. WT when AA results were included or excluded, respectively. ns = not significant. N=7/group. D: 4-6 h fasting plasma insulin levels vs. 4-6 h fasting blood glucose levels for each mouse whose data is represented in (A). Data were fit via linear regression. The significance of the slope deviating from zero (p-value) and goodness-of-fit (R^2^ value) for each dataset are shown. E: Plasma insulin values in (D) normalized to their own matching blood glucose values. Data represent mean ± SEM and were compared by two-way T test. * or †, p < 0.05 vs. WT when AA results were included or excluded, respectively. N=7/group.

While the findings in Figures 5D and 5E are supportive of improved β-cell function in Gα_z_KO NOD mice, as a strain characteristic, NOD mice are extremely insulin sensitive (as reflected in the weak stimulation of plasma insulin levels after oral glucose challenge), making *in vivo* determination of β-cell function challenging. Therefore, to directly assess β-cell function, islets were isolated from control diet or EPA diet-fed male and female NOD mice and used in *ex vivo* GSIS assays in low glucose (1.7 mM; ~30 mg/dl) or stimulatory glucose (16.7 mM; ~300 mg/dl), the latter with or without 10 nM sulprostone: an EP3-selective agonist. While the mean islet insulin content of WT control diet-fed mice was lower than those of the other groups, this differences was not statistically-significant (**Figure 6A**). Islets from Gα_z_KO mice secreted more insulin as a percent of content in stimulatory glucose as compared to WT, with the insulin secretion response of EPA diet-fed Gα_z_KO mice being significantly elevated over both WT control diet-fed and WT EPA-diet fed. (**Figure 6B**). Sulprostone had no significant effect on GSIS in islets from any of the groups (**Figure 6C**). Quantitative PCR using primers specific for mRNA encoding each of the three mouse EP3 splice variants, EP3α, EP3β, and EP3*γ*, reveals all are significantly elevated in islets from 16-17-week-old NOD mice as compared to 10-12-week old C57BL/6N mice (**Figure 6D**): a strain which has no *in vivo* glucose tolerance or *in vivo* or *ex vivo* insulin secretion phenotype of the Gα_z_ knockout (4,6).

**Figure 6.**
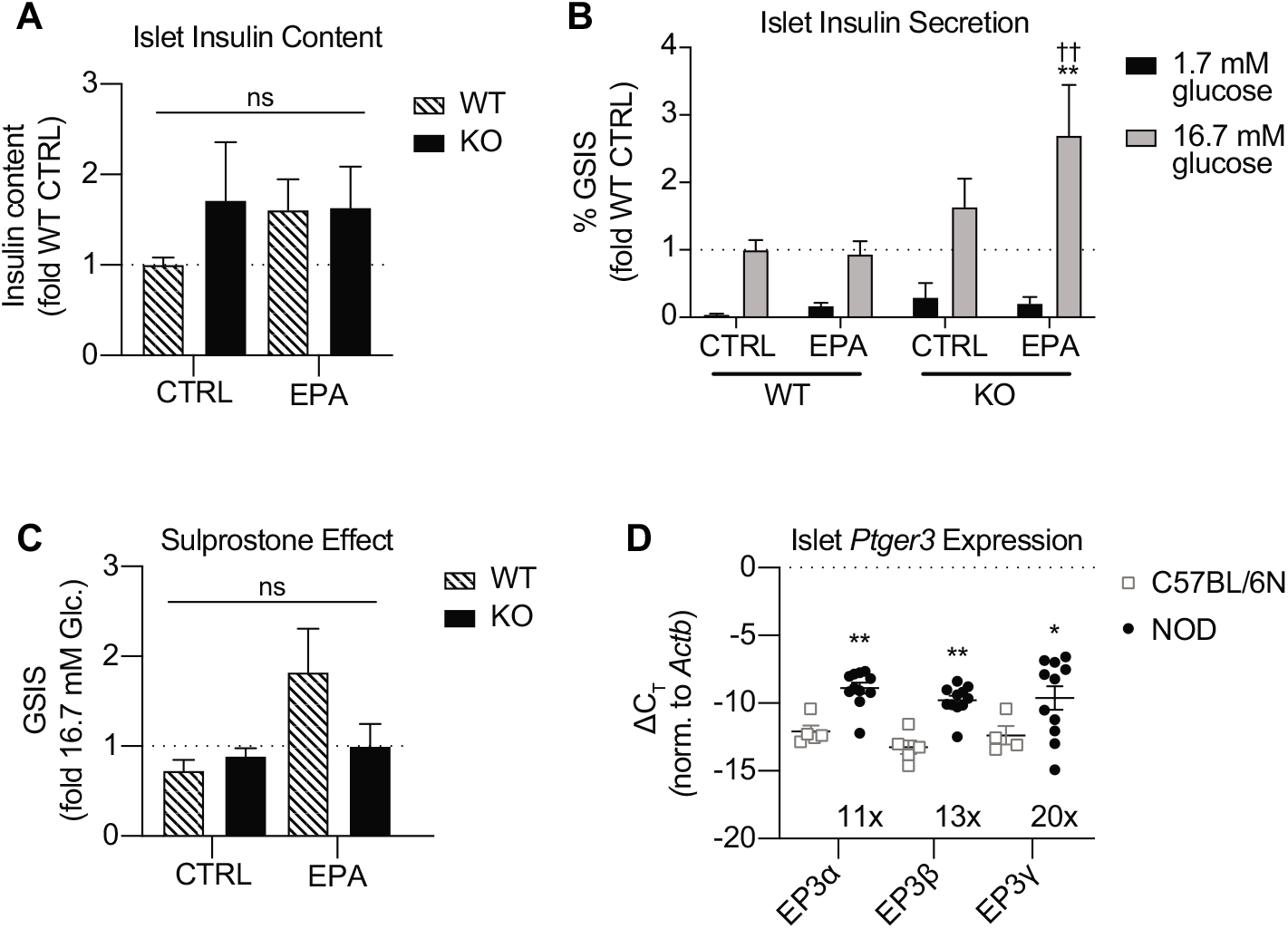
An EPA-enriched diet enhances NOD islet glucose stimulated insulin secretion (GSIS) only when Gα_z_ is lost, independent of agonist-sensitive EP3 activity. A-C: (A) Islet insulin content, (B) insulin secreted as a percent of content in 1.7 mM glucose (~30 mg/dl; nonstimulatory) or 16.7 mM (~300 mg/dl; stimulatory), and (D) the effect of 10 nM sulprostone on 16.7 mM GSIS in islets from male and female NOD mice fed the control diet or EPA-enriched diet for 12 weeks upon weaning. In all panels, data represent mean ± SEM and data were compared by two-way ANOVA with Holm-Sidak test post-hoc to correct for multiple comparisons. In (B), a paired analysis was performed. **, p < 0.01 vs. WT Control and ††, p < 0.01 vs. WT EPA. N=7-9/group. D: Results of quantitative PCR (qPCR) analysis of male and female NOD islet cDNA samples using primers specific for each of the three mouse EP3 splice variants, EP3α, EP3β, and EP3*γ*, as compared to islet cDNA samples from 10-12-week old male C57BL/6N mice (described in (6)). Data represent mean ± SEM, and means were used to calculate relative changes in abundance between mouse strains, as indicated, via the 2^ΔΔC^_T_ method. Data were compared by two-way ANOVA with Holm-Sidak test post-hoc to correct for multiple comparisons. *, p < 0.05 and **, p < 0.01 vs. C57BL/6N. N=5-10 per group.

## Discussion

Islet PGE_2_ production, circulating PGE_2_ metabolite levels, and/or islet EP3 expression are well-established as increased in mice and humans with T2D (4,9,10,14,19–24), with evidence of this mechanism being conserved in patients with T1D or mouse models of the disease (25–29). Gα_z_ is critical for mediating β-cellEP3 signaling (4–7). Gα_z_KO NOD mice are protected from developing hyperglycemia, and NOD mice fed a diet enriched in EPA have improved glucose tolerance (8,9). In the past, this led us to propose a dietary intervention to reduce PGE_2_ production or a therapeutic designed to interfere with β-cellEP3/ Gα_z_ interaction could both be valid strategies to improve the β-cell dysfunction of T1D, perhaps even providing a synergistic effect (11,19). This hypothesis was tested in the current work, with surprising results.

Pivotal findings on the roles of PUFAs in β-cell function, particularly in the context of T1D, have come out of studies with the mFat-1 model, in which mice transgenically express the *C. elegans* n-3 fatty acid desaturase, *fat-1*, rendering the mouse capable of converting omega-6 to omega-3 PUFAs and shifting omega-6-to-omega-3 PUFA ratios (30). Wei and colleagues showed islets from mFat-1 mice had improved islet insulin secretion and decreased β-cell death after cytokine treatment (31). Wang and colleagues showed mFat-1 mice were resistant to streptozotocin-induced T1D, and that EPA and DHA, but not AA, prevented tunicamycin induced β-cell ER stress *ex vivo* (32). Complementary to these findings, Bi and colleagues showed an omega-3 supplemented diet significantly improved the NOD T1D phenotype (33). A major caveat to these previous studies, though, is omega-3 PUFAs were given at artificially high levels, whether endogenously produced or exogenously administered. Our study employed a diet with physiologically-relevant PUFA concentrations, validated to have the expected impact on circulating PGE_2_ metabolite levels in NOD mice, regardless of genotype. In contrast to our expectations, the glucose tolerance of both AA diet- and EPA diet-fed NOD mice improved as compared to control diet-fed WT animals, suggesting a phenotype independent of PGE_2_.

Islet inflammation in the context of T1D is mediated by a host of different innate and adaptive immune cells. T-cell activation and recruitment to the islet is known to facilitate recognition of auto-antigens and islet destruction. An imbalance of Th17 and Tregs has been found in T1D (34). Restoration of Treg activity has been shown to ameliorate the NOD T1D phenotype, and Treg function is the target of developing T1D therapeutics (35–37). High-dose omega-3 PUFA supplementation has previously been shown to alter splenic T-cell populations (33). In the current work, a physiologically-relevant EPA-supplemented diet significantly increased the abundance of Th1 and Treg T cells in 29-30-week old NOD females, with no effect on Th17 T-cells. Yet, the caveat to this and previous studies is only the effects of omega-3 and/or EPA supplementation were determined, not AA.

In support of a general PUFA effect on the NOD immune phenotype, IL-11 mRNA expression was increased more than 100-fold with either AA diet feeding or EPA diet feeding in islets from WT mice. IL-11, an anti-inflammatory cytokine, inhibits NFκB activation and downstream production of proinflammatory cytokines such as TNF, IL-1β, and IFNɣ, specifically in the context of T1D (38–40). PUFA-induced alterations in splenic T-cell populations and islet cytokine expression, though, did not translate to changes in islet insulitis profile or insulin-positive pancreas area. Yet, this does not exclude a real effect of PUFAs in the maintenance of functional β-cell mass. In support of this concept, CXCL10 recruits activated T-cells to the islet, and neutralization of CXCL10 can enhance β-cell proliferation without impacting insulitis (41). Furthermore, plasma levels of EPA have been associated with improved fasting C-peptide concentration, a read-out of residual functional β-cell mass, in youth with recent onset T1D (42).

While Gα_z_ loss had no effect on splenic T-cell populations, confirmatory of previous work (8), Gα_z_ loss on its own was associated with improved islet cytokine expression, insulitis profile, β-cell fractional area, and β-cell function, with, in some cases, a PUFA-enriched diet moderating these changes. Our original model that enriching islets with EPA, thereby reducing the substrate for PGE_2_ production, would coordinate with Gα_z_ loss to improve β-cell function was indeed borne out in islet GSIS assays. Yet, neither WT nor Gα_z_KO islets responded to the EP3 agonist, sulprostone, to reduce GSIS. Compared to islets from young, lean C57BL/6N mice, in which no *in vivo* or *ex vivo* islet phenotype of Gα_z_ loss exists (4,6), the expression of all three EP3 splice variants in NOD islets is 10-20-fold enhanced. Notably, EP3α and EP3*γ* have partial-to nearly-full agonist-insensitive activity, respectively (43–45) and islet EP3*γ* signals exclusively through Gα_z_ (14). Future mechanistic studies on the role of AA- and EPA supplementation on the systemic and islet immune phenotype in T1D and how this coordinates with EP3/ Gα_z_-mediated regulation of β-cell function and mass are critically important areas for future investigation.

## Acknowledgements

We wish to thank the many present and former members of the Kimple Laboratory who contributed technical assistance or scientific discussion during the course of these experiments, particularly Joshua Neuman, who designed and characterized the diets used in this work. This work was supported in part by Merit Review Award I01 BX003700 from the United States (U.S.) Department of Veterans Affairs Biomedical Laboratory Research and Development Service (BLR&D) (to MEK). Further support was provided by NIH Grants R01 DK102598 (to MEK), F31 DK109698 (to RJF), and R01 DK109508 (to JG); ADA Grant 1-16-IBS-212 (to MEK), and JDRF Grant 17-2011-608 (to MEK). MEK is the guarantor of this work and, as such, had full access to all the data in the study and takes responsibility for the integrity of the data and the accuracy of the data analysis.

## Author Contributions

Conceptualization, RJF, JG, and MEK; data curation, RJF and MEK; formal analysis, RJF, DCP, MDS, AP, and MEK; funding acquisition, RJF, JG, and MEK; investigation, RJF, HNW, DCP, MDS, and AP; methodology, RJF, AP, JG, and MEK; project administration, MEK; supervision, RJF, JG, and MEK; visualization, RJF, AP, and MEK; writing—original draft, RJF and DCP; writing—review and editing, RJF, DCP, AP, and MEK. All authors have read and agreed to the published version of the manuscript.

